# Host-directed therapies against early-lineage SARS-CoV-2 retain efficacy against B.1.1.7 variant

**DOI:** 10.1101/2021.01.24.427991

**Authors:** Ann-Kathrin Reuschl, Lucy G. Thorne, Lorena Zuliani-Alvarez, Mehdi Bouhaddou, Kirsten Obernier, Joseph Hiatt, Margaret Soucheray, Jane Turner, Jacqueline M. Fabius, Gina T. Nguyen, Danielle L. Swaney, Romel Rosales, Kris M. White, Pablo Avilés, Ilsa T. Kirby, James E. Melnyk, Ying Shi, Ziyang Zhang, Kevan M. Shokat, Adolfo García-Sastre, Clare Jolly, Gregory J. Towers, Nevan J. Krogan

## Abstract

Coronavirus disease 2019 (COVID-19), caused by severe acute respiratory syndrome coronavirus 2 (SARS-CoV-2), has resulted in millions of deaths worldwide and massive societal and economic burden. Recently, a new variant of SARS-CoV-2, known as B.1.1.7, was first detected in the United Kingdom and is spreading in several other countries, heightening public health concern and raising questions as to the resulting effectiveness of vaccines and therapeutic interventions. We and others previously identified host-directed therapies with antiviral efficacy against SARS-CoV-2 infection. Less prone to the development of therapy resistance, host-directed drugs represent promising therapeutic options to combat emerging viral variants as host genes possess a lower propensity to mutate compared to viral genes. Here, in the first study of the full-length B.1.1.7 variant virus, we find two host-directed drugs, plitidepsin (aplidin; inhibits translation elongation factor eEF1A) and ralimetinib (inhibits p38 MAP kinase cascade), as well as remdesivir, to possess similar antiviral activity against both the early-lineage SARS-CoV-2 and the B.1.1.7 variant, evaluated in both human gastrointestinal and lung epithelial cell lines. We find that plitidepsin is over an order of magnitude more potent than remdesivir against both viruses. These results highlight the importance of continued development of host-directed therapeutics to combat current and future coronavirus variant outbreaks.

## Introduction

The severe acute respiratory syndrome coronavirus 2 (SARS-CoV-2), the causative agent of coronavirus disease 2019 (COVID-19), has spread throughout the world causing millions of deaths and massive societal and economic disruption. In September 2020, a novel variant of SARS-CoV-2 was detected in the United Kingdom (UK), known as lineage B.1.1.7. This variant has rapidly spread since November 2020 around the UK and now comprises up to 40% of COVID-19 new cases in the South East of England (Public Health England, 3rd technical briefing). Critically, B.1.1.7 has also been detected in numerous countries around the world, including the United States, Europe, and Canada. Rapidly increasing numbers of B.1.1.7 infections and B.1.1.7 displacement of earlier lineages in the UK exemplify constant changes in circulating viral strains and intensify the challenge of containing viral spread.

Sequencing of the B.1.1.7 variant revealed 23 mutations (Rambaut et al., 2020); 17 protein-coding mutations (14 non-synonymous mutations and 3 deletions) and 6 synonymous mutations (Figure 1). The majority of the protein-coding mutations (8) were observed within the viral Spike protein, which facilitates viral entry through its interaction with the human ACE2 receptor. Recent reports have suggested that changes in Spike, especially in the receptor binding domain (RBD), found in new SARS-CoV-2 variants may alter antibody neutralization sensitivity and possibly vaccine efficacy (Collier et al., 2021; Reese-Spear et. al, 2021), though this does not appear to be true for Spike RBD mutation N501Y of the B.1.1.7 variant (Muik et al., 2021; Rathnasinghe et al, 2021). The other non-synonymous mutations are located in Nsp3 (3), Nsp6 (1), Orf8 (3), and N (2) protein. The impact of these mutations on viral replication, transmission and pathogenesis are still not well understood. It is paramount to investigate the impact of these mutations on the viral life cycle and evaluate the efficacy of previously identified candidate antiviral treatments for COVID-19 on this new SARS-CoV-2 variant.

**Figure 1.**
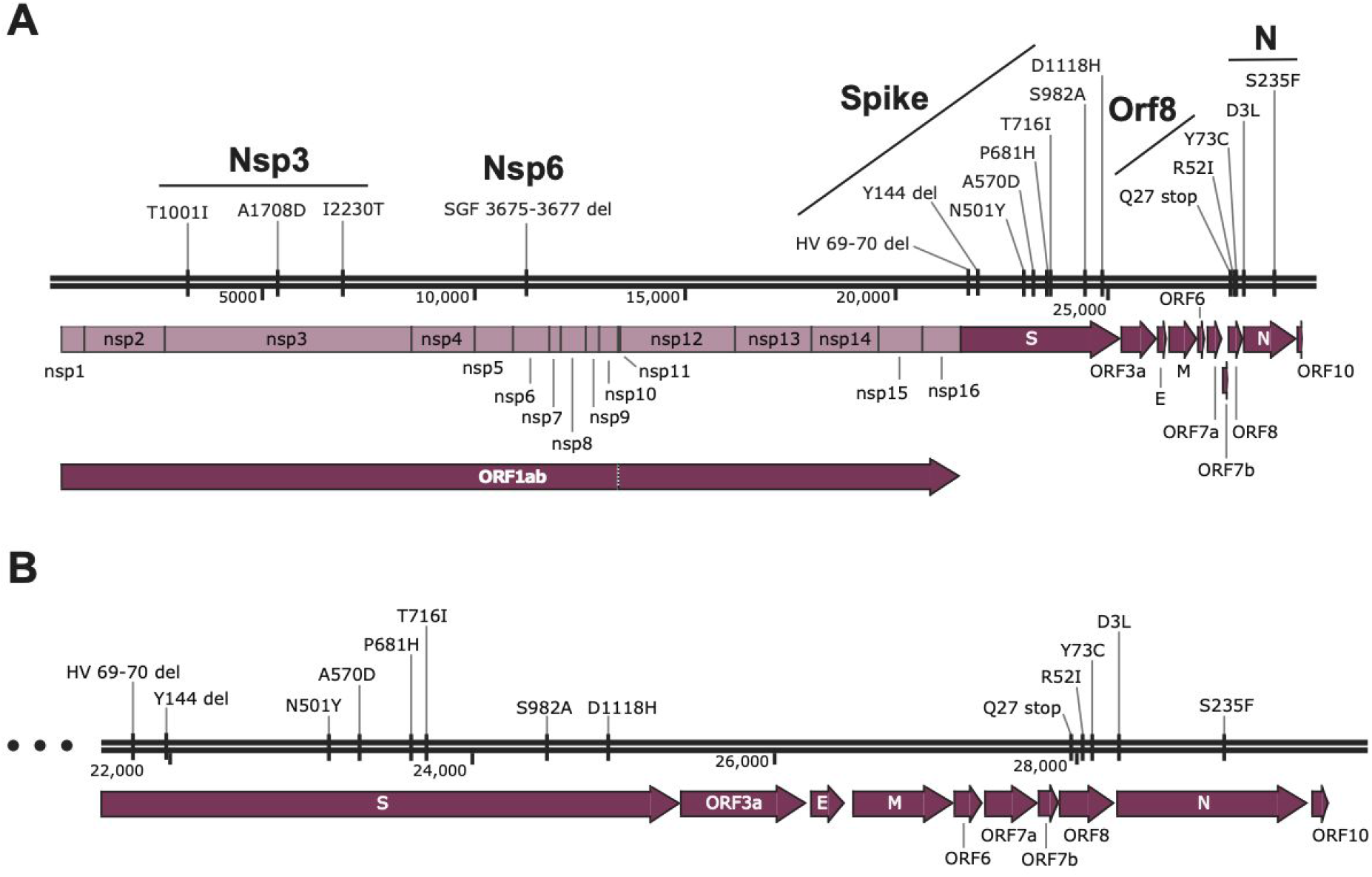
Protein-coding mutations in SARS-CoV-2 B.1.1.7 virus variant. **A)** SARS-CoV-2 genome map (NC_045512.2) depicting open reading frames (dark purple arrows) and proteins resulting from cleavage of ORF1ab polypeptide into non-structural proteins (light purple). Seventeen protein-coding mutations are annotated (top) which includes 14 non-synonymous mutations and 3 deletions spanning 5 viral proteins. **B)** Expansion of the genomic region from spike to N protein for visualization purposes.

Currently, only immunosuppressive interventions such as dexamethasone have been successful in reducing mortality in severe COVID-19 cases (RECOVERY Collaborative Group et al., 2020). There remains an urgent need to test and develop new therapeutic interventions for clinical treatment that target both viral replication and immunopathology associated with poor prognosis. Host-directed therapies offer advantages over those directly targeting the virus, as host genes possess a lower propensity to mutate, and present a higher barrier to viral adaptation involving interactions with essential host co-factors. Moreover, druggable host targets often play roles in other diseases, providing the opportunity to repurpose FDA-approved drugs and expedite drug development timelines.

Previously, we and others have identified several host factors that directly interact with SARS-CoV-2 proteins (Gordon, Hiatt, et al., 2020; Gordon, Jang, et al., 2020), or are impacted during the course of viral infection (Bouhaddou et al., 2020). Many of these host factors are targeted by FDA-approved drugs, investigational new drugs (INDs), or preclinical compounds, and demonstrated strong antiviral effects. For example, we uncovered that the eukaryotic translation factor eIF4H interacts with SARS-CoV-2 Nsp9 and identified inhibitors of the eukaryotic translation machinery as potent antivirals against SARS-CoV-2, including zotatafin (inhibitor of eIF4A) and ternatin-4 (inhibitor of eEF1A), a molecular derivative of plitidepsin (aplidin) (Gordon, Jang, et al., 2020). Plitidepsin, another eEF1A inhibitor clinically approved in some countries to treat multiple myeloma (Leisch et al., 2019), possessed potent antiviral activity in several cell lines, primary cells, and mouse models (White et al., 2021). Furthermore, pharmacological inhibition of p38 signaling using ralimetinib and other p38 inhibitors demonstrated strong antiviral activity, consistent with a marked increase in p38-MAP kinase activity during SARS-CoV-2 infection (Bouhaddou et al., 2020). Here, we show that plitidepsin, ralimetinib, and remdesivir (a widely-used broad-spectrum antiviral nucleoside analog approved for use in several countries for treatment of COVID-19) possess antiviral efficacy against the B.1.1.7 virus variant.

## Results/Discussion

We first treated intestinal epithelial Caco-2 cells with a range of concentrations of plitidepsin, ralimetinib, or remdesivir prior to infection with SARS-CoV-2 early-lineage BetaCoV/Australia/VIC01/2020 (VIC) or the B.1.1.7 variant (SARS-CoV-2 England/ATACCC 174/2020 isolate) for 24 hours. Plitidepsin potently suppressed viral replication of both SARS-CoV-2 strains in the nanomolar range, measured by intracellular nucleocapsid protein positive cells (% N+ cells) by flow cytometry (Fig. 2 A) as well as viral RNA replication measured by accumulation of genomic and subgenomic E RNA by RT-qPCR (Fig. 2 B). Inhibition of viral replication was also observed with the p38-targeting inhibitor ralimetinib, with antiviral effects manifesting at µM drug concentrations (Fig. 2 D and E). We observed no cytotoxicity over the range of concentrations used for each inhibitor by MTT assay (Fig. 2 C and F). Remdesivir, a polymerase inhibitor (Kovic, 2020), was used as a positive control and also inhibited replication of both viruses in a dose-dependent manner in the µM range (Fig.2H), as observed elsewhere (Thorne et al., 2020), with no observed cytotoxicity (Fig. 2 I). Strikingly, plitidepsin was effective against the early and more recent SARS-CoV-2 lineages at concentrations over an order of magnitude lower than remdesivir.

**Figure 2.**
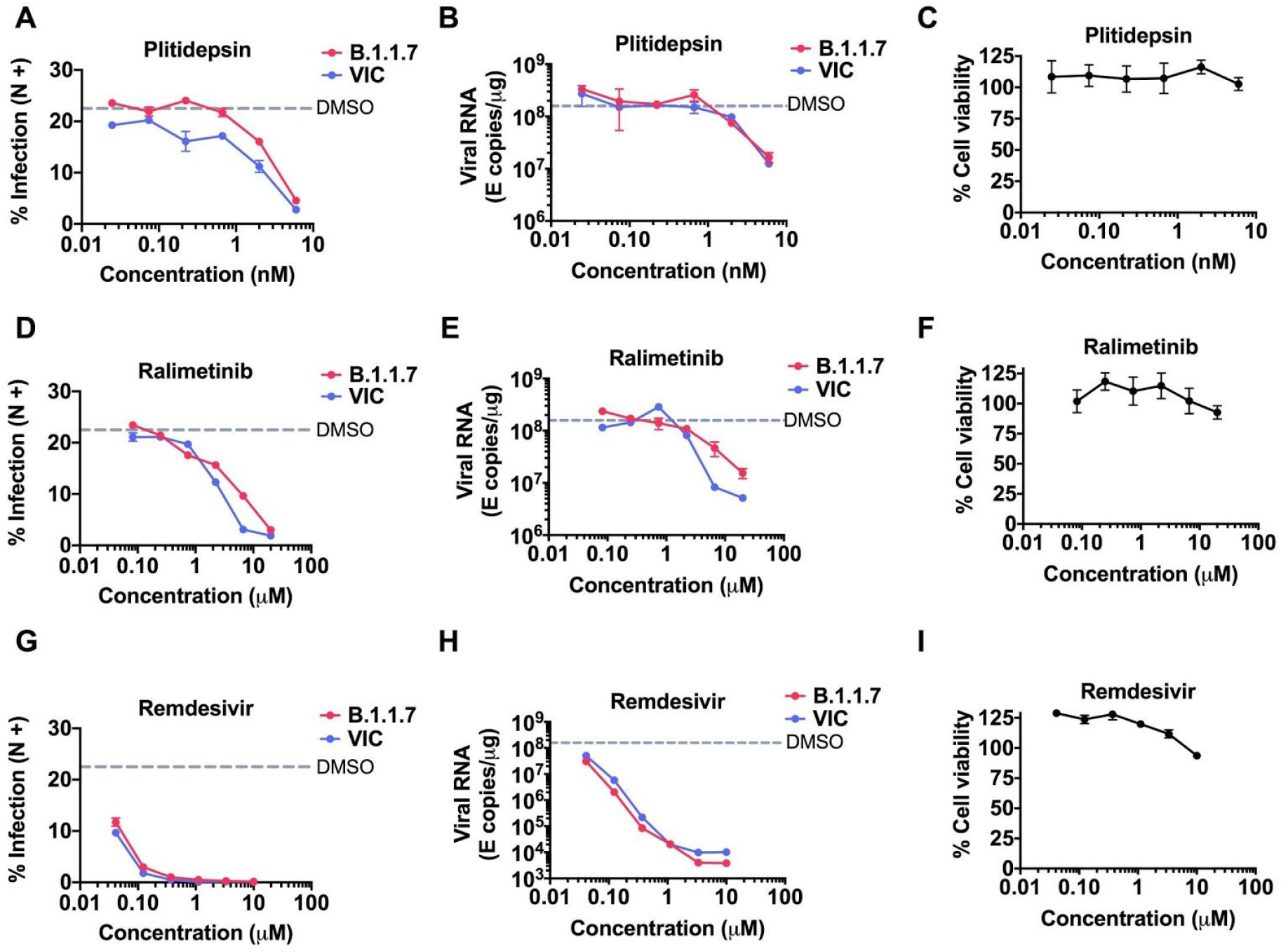
Antiviral efficacy against early-lineage and B.1.1.7 variant SARS-CoV-2 in Caco-2 human intestinal epithelial cells. BetaCoV/Australia/VIC01/2020 (VIC) or SARS-CoV-2 B.1.1.7 (SARS CoV 2 England/ATACCC 174/2020) replication after treatment with increasing doses of eEF1A-inhibitor plitidepsin, p38-inhibitor ralimetinib or viral replication inhibitor remdesivir. (**A, D, G**) Infection levels in the presence of plitidepsin (A), ralimetinib (D) or remdesivir (G) were measured by intracellular staining for SARS-CoV-2 nucleocapsid protein (N+) at 24h post infection. (**B, E, H**) Replication of SARS-CoV-2 genomic and subgenomic E RNAs per μg total RNA measured by qRT-PCR is shown. (**C, F, I**) Cell viability in the absence of infection after exposure to indicated inhibitor concentrations is shown relative to DMSO treated controls at 24 hours. Dashed lines indicate viral replication in the presence of DMSO vehicle control. Mean+/-SEM (n=3).

To confirm that both host-targeted drugs retained activities in innate-sensing competent cells, we tested their effect on SARS-CoV-2 infection in Calu-3 human lung epithelial cells. This cell line is highly permissible to SARS-CoV-2 infection, while mounting a robust inflammatory innate immune response (Thorne et al 2020) similar to that observed in primary human airway cells (Ravindra et al. 2020). We tested the highest non-toxic concentration (>75% cell viability) of each drug (Fig. 3 A, D, G), 6nM plitidepsin, 6.7µM ralimetinib and 1.1µM remdesivir, against both SARS-CoV-2 lineages. Similar to what we observed in Caco-2 cells, plitidepsin retained the most potent antiviral effects against B.1.1.7 and BetaCoV/Australia/VIC01/2020 (VIC) at nM concentration (Fig. 3 B and C), but all three drugs effectively suppressed replication of both viruses (Fig.3 B,C,E,F,H,I).

**Figure 3.**
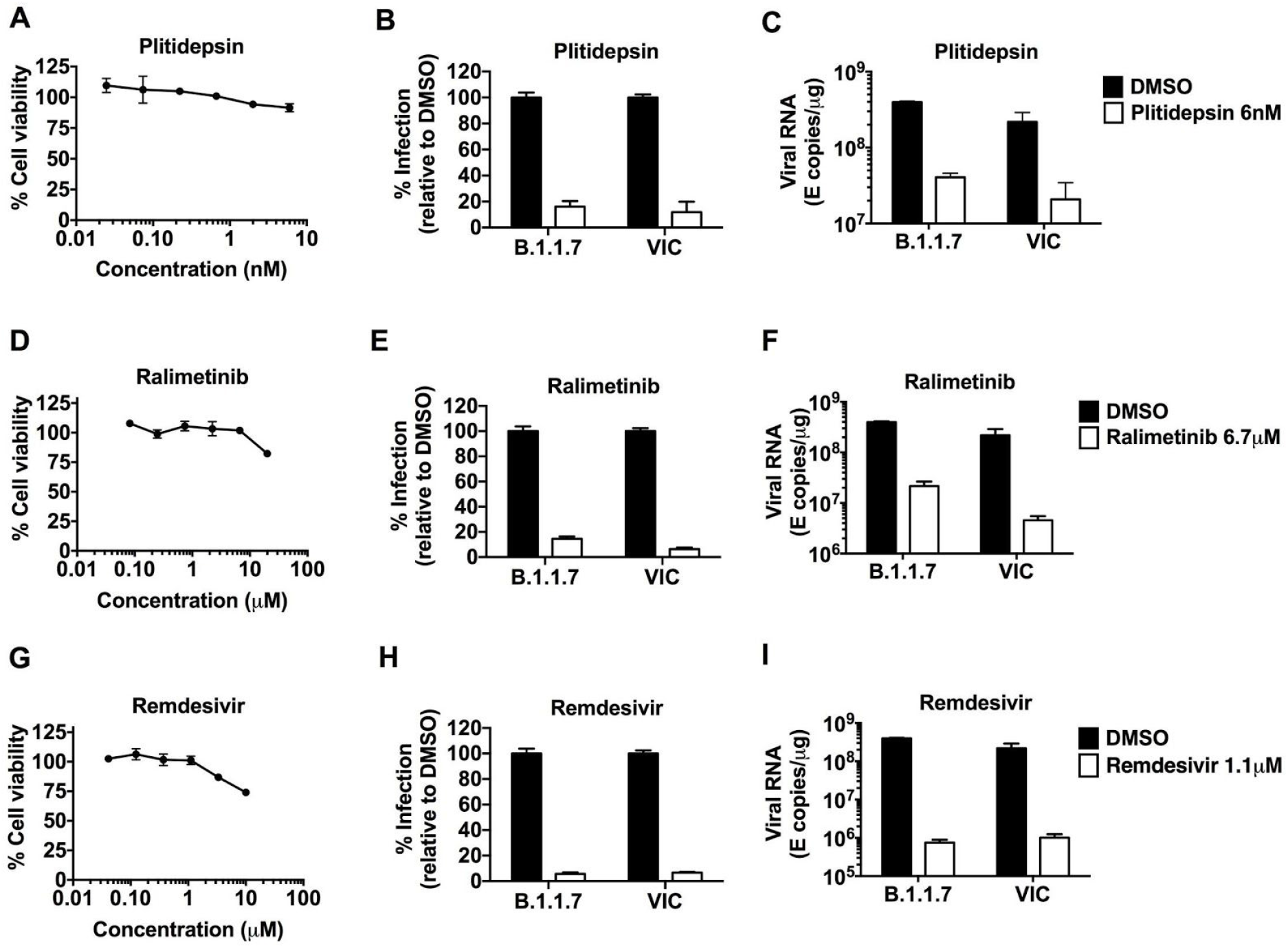
Antiviral efficacy against early-lineage and B.1.1.7 variant SARS-CoV-2 in Calu-3 human lung epithelial cells. Viral replication in Calu-3 human lung epithelial cells of SARS-CoV-2 BetaCoV/Australia/VIC01/2020 (VIC) or SARS-CoV-2 B.1.1.7 (SARS CoV 2 England/ATACCC 174/2020) after treatment with increasing doses of eEF1A-inhibitor plitidepsin, p38-inhibitor ralimetinib or viral replication inhibitor remdesivir (n=3). (**A, D, G**) Cell viability in the absence of infection after exposure to indicated inhibitor concentrations is shown relative to DMSO treated controls at 24 hours. (**B, E, H**) Infection levels after treatment with plitidepsin (6nM) (B), ralimetinib (6.7 μM) (E) or remdesivir (1,1 μM) (H) were measured by intracellular staining for SARS-CoV-2 nucleocapsid protein (N+) at 24h post infection. Percentage infection is shown relative to DMSO-treated controls (n=2-3). (**C, F, I**) Replication of SARS-CoV-2 genomic and subgenomic E RNAs per μg total RNA measured by qRT-PCR is shown. Dashed lines indicate viral replication in the presence of DMSO vehicle control. Mean +/-SEM (n=3).

Taken together, the host-directed drugs plitidepsin and ralimetinib, as well as remdesivir possess antiviral efficacy against both the SARS-CoV-2 early-lineage (VIC) and the B.1.1.7 virus variant. Plitidepsin is a clinical-approved therapeutic for multiple myeloma (Leisch et al., 2019) and is currently in a phase 2 clinical trial for treatment of COVID-19 in Spain (https://clinicaltrials.gov/ct2/show/NCT04382066), due to its antiviral properties in cell culture and animal models of disease. Consistent with the data above, we have recently shown that plitidepsin is 27.5-fold more potent against SARS-CoV-2 early-lineage than remdesivir and exerts antiviral activity with an IC50 of approximately 1 nM (White et al., 2021). Plitidepsin is an antagonist of eEF1A, which is part of the eukaryotic translation elongation 1 complex and commonly used by RNA viruses for mRNA translation (Li et al., 2013). Therefore, treatment with plitidepsin can successfully interfere with viral replication and might be extended to other viral infections as a broad-acting antiviral, which would be consistent with previous reports suggesting that viruses are more susceptible to regulators of proteostasis than their cellular hosts (Heaton et al., 2016).

Ralimetinib, a p38 mitogen-activated protein kinase inhibitor, was initially developed for the treatment of patients with advanced cancer (Patnaik et al., 2016; Vergote et al., 2020). p38 is involved in different cellular processes including protein translation, cell cycle, inflammation and cell death (Zarubin & Jiahuai, 2005). We previously reported that p38-MAPK is strongly upregulated during SARS-CoV-2 infection and drugs targeting these pathways showed antiviral activity *in vitro*, suggesting that SARS-CoV-2 relies on p38 activation for viral replication. Similar findings have been reported for SARS-CoV and HCoV-229E, where activation of p38 is observed during viral infection and inhibition of this pathway can limit viral replication (Kono et al., 2008; Mizutani et al., 2004). Moreover, we have demonstrated that blocking p38 activation significantly reduces cytokine expression (Bouhaddou et al., 2020), suggesting that targeting this pathway can be beneficial in severe cases of COVID-19, where immune dysregulation and tissue damage are associated with poor disease outcome.

While mutations in the Spike protein in the B.1.1.7 variant could potentially contribute to increased transmissibility, mutations in other viral proteins could also be playing a role. It will be interesting to determine if and how these mutations alter the SARS-CoV-2-human protein-protein interaction landscape (Gordon, Hiatt, et al., 2020; Gordon, Jang, et al., 2020). Also, analysis of cellular changes in the host during infection with the different lineages, including monitoring alterations in signaling networks via post-translational modifications (i.e. phosphorylation (Bouhaddou et al., 2020) and ubiquitination) as well as host genetic dependencies (Daniloski et al., 2021; Wang et al., 2021; Wei et al., 2021), may help to understand these and future variants.

This study builds on our prior characterization of host-directed drugs as antivirals for SARS-CoV-2 infection (Gordon, Hiatt, et al., 2020; Gordon, Jang, et al., 2020; Bouhaddou et al., 2020; White et al., 2021) and demonstrate the advantages of repurposing drugs that are clinically approved with known pharmacokinetics and safety profiles, as they can quickly transition into clinical trials. Furthermore, our data encourage the continued pre-clinical and clinical development of host-directed antiviral therapies for COVID-19, as targeting essential host-factors required for viral replication provides a strategy to retain efficacy against emerging SARS-CoV-2 variants. Importantly, as these drugs inhibit cellular processes commonly employed by different viruses to ensure infection, these and other host-directed therapies could be used to rapidly respond to future viral outbreaks.

## Acknowledgments

This research was funded by grants from the National Institutes of Health (P50AI150476, U19AI135990, U19AI135972, R01AI143292, R01AI120694, P01A1063302, and R01AI122747 to N.J.K. and F32CA239333 to M.B.), Defense Advanced Research Projects Agency (DARPA) (#HR0011-19-2-0020 to K.M.S, N.J.K., and A.G.-S.), by the Excellence in Research Award (ERA) from the Laboratory for Genomics Research (LGR), a collaboration between UCSF, UCB, and GSK (#133122P to N.J.K.), by a Fast Grant for COVID-19 from the Emergent Ventures program at the Mercatus Center of George Mason University to N.J.K., by the Roddenberry Foundation to N.J.K., by funding from F. Hoffmann-La Roche and Vir Biotechnology and gifts from QCRG philanthropic donors. J.H. was supported by the UCSF MSTP (T32GM007618). NIH grant 1F32CA236347-01 (to J.E.M.); Z.Z. is a Damon Runyon Fellow supported by the Damon Runyon Cancer Research Foundation (DRG-2281-17); K.M.S. acknowledges support from HHMI. G.J.T. is funded by a Wellcome Senior Fellowship. This research was also partly funded by by CRIP (Center for Research for Influenza Pathogenesis), a NIAID supported Center of Excellence for Influenza Research and Surveillance (CEIRS, contract # HHSN272201400008C); by NCI grant U54CA260560; by supplements to NIAID grant U19AI135972 and DoD grant W81XWH-20-1-0270; by the generous support of the JPB Foundation and the Open Philanthropy Project (research grant 2020-215611 (5384); and by anonymous donors to A.G-S. C.J. is funded by a Welcome Investigator award. Funds were also obtained from the University College London COVID-19 fund and the National Institutes of Health Research UCL/UCLH Biomedical Research Centre. We are grateful to the National Institute of Health Research Health Protection Research Unit in Respiratory Infections (NIHR #200927), the Assessment of Transmission and Contagiousness of COVID-19 in Contacts (ATACCC) Study funded by the DHSC COVID-19 Fighting Fund and the ATACCC investigators, in particular Joe Fenn, Rhia Kundu, Robert Varro, Sarah Hammett, Jessica Cutajaar, Eimear McDermott, Jada Samuel, Samuel Bremang, Alexandra Koycheva, Nieves Fernandez Derqui, Sam Janakan, Emily Conibear, Lulu Wang & Seran Hakki. We are also grateful to Ajit Lalvani, Jake Dunning, Maria Zambon and colleagues at Public Health England and Giada Mattiuzzo at the National Institute for Biological Standards and Controls and Wendy Barclay and Jonathan Brown and all colleagues in the United Kingdom Research Institute funded collaboration Genotype to Phenotype for facilitating our obtaining SARS-CoV-2 isolates. We are grateful to Laura McCoy at UCL for providing the SARS2 CoV-2 N antibody.

## Methods

### Cells culture and drugs

Calu-3 cells (ATCC HTB-55) and Caco-2 cells were a kind gift Dr Dalan Bailey (Pirbright Institute). VeroE6 cells were provided by NIBSC. Cells were cultured in Dulbecco’s modified Eagle Medium (DMEM) supplemented with 10% heat-inactivated FBS (Labtech), 100U/ml penicillin/streptomycin, with the addition of 1% Sodium Pyruvate (Gibco) and 1% Glutamax for Calu-3 and Caco-2 cells. All cells were passaged at 80% confluence. For infections, adherent cells were trypsinized, washed once in fresh medium and passed through a 70 µm cell strainer before seeding at 0.2×10^6 cells/ml into tissue-culture plates. Calu-3 cells were grown to 60-80% confluence prior to infection as described previously (Thorne et al 2020). Plitidepsin (aplidin) (PharmaMar), ralimetinib (SelleckChem), remdesivir (SelleckChem) were reconstituted in sterile DMSO.

### Viruses

SARS-CoV-2 strain BetaCoV/Australia/VIC01/2020 (NIBSC) and SARS-CoV-2 B.1.1.7 (SARS CoV 2 England/ATACCC 174/2020) strain were propagated by infecting Caco-2 cells at MOI 0.01 TCID50/cell, in DMEM supplemented with 2% FBS at 37°C. Virus was harvested at 72 hours post infection (hpi) and clarified by centrifugation at 4000 rpm for 15 min at 4 °C to remove any cellular debris. Virus stocks were aliquoted and stored at -80 °C. Virus titres were determined by 50% tissue culture infectious dose (TCID50) on Vero.E6 cells. In brief, 96 well plates were seeded at 1×10^4 cells/well in 100 µl. Eight ten-fold serial dilutions of each virus stock or supernatant were prepared and 50 µl added to 4 replicate wells. Cytopathic effect (CPE) was scored at 5 days post infection, and TCID50/ml was calculated using the Reed & Muench method (Reed & Muench, 1938), and an Excel spreadsheet created by Dr. Brett D. Lindenbach was used for calculating TCID50/mL values (Lindenbach, 2009).

### Infection and drug assays

Calu-3 and Caco-2 cells were pre-treated with remdesivir, plitidepsin or ralimetinib at the indicated concentrations or DMSO control at an equivalent dilution for 2 h before SARS-CoV-2 infection. Caco-2 cells were infected at 4.8×10^3 E copies per cell, equivalent to an MOI of 0.04 TCID50 per cell (as titred on Vero.E6), and Calu-3 cells were infected at 1.2×10^3 E copies per cell, equivalent to an MOI of 0.01 TCID50 per cell, in the presence of the indicated inhibitor concentrations, and incubated for 2h at 37 °C. After 2h, the inoculum was removed and fresh culture medium was added containing the inhibitors, which were maintained throughout infection. Cells were harvested after 24h for analysis and viral infection measured by quantification of E genomic and subgenomic RNA copies by RT-qPCR and intracellular detection of SARS-CoV-2 nucleoprotein by flow cytometry. Tetrazolium salt (MTT) assay was performed to verify cell viability. 10 % v/v MTT was added to the cell media and cells were incubated for 2 h at 37°C. Cells were lysed with 10% SDS, 0.01M HCl and the formation of purple formazan was measured at 620nm.

### RT-qPCR

RNA was extracted using RNeasy Micro Kits (Qiagen) and residual genomic DNA was removed from RNA samples by on-column DNAse I treatment (Qiagen). Both steps were performed according to the manufacturer’s instructions. cDNA was synthesized using SuperScript III with random hexamer primers (Invitrogen). RT-qPCR was performed using Fast SYBR Green Master Mix (Thermo Fisher) for host gene expression or TaqMan Master mix (Thermo Fisher) for viral RNA quantification, and reactions performed on the QuantStudio 5 Real-Time PCR systems (Thermo Fisher). Viral RNA copies were deduced by a standard curve, using primers and a Taqman probe specific for E, as described elsewhere (Corman et al., 2020) and below. The following primers and probes were used:

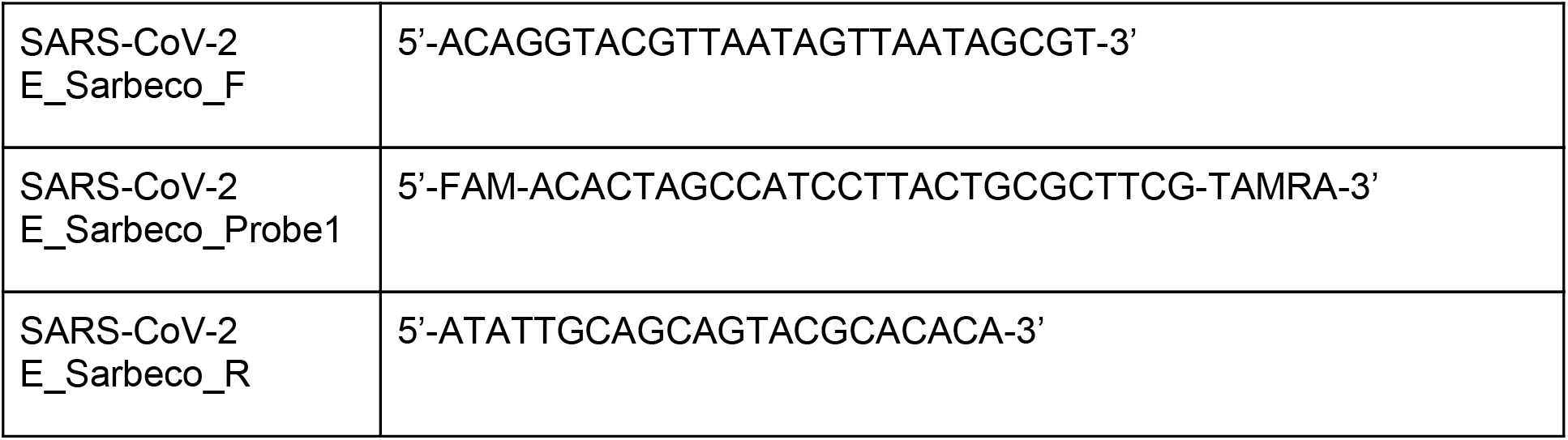

### Flow cytometry

For flow cytometry analysis, adherent cells were recovered by trypsinization and washed in PBS with 2mM EDTA (PBS/EDTA). Cells were stained with fixable Zombie UV Live/Dead dye (Biolegend) for 6 min at room temperature. Excess stain was quenched with FBS-complemented DMEM. Unbound antibody was washed off thoroughly and cells were fixed in 4% PFA prior to intracellular staining. For intracellular detection of SARS-CoV-2 nucleoprotein, cells were permeabilized for 15 min with Intracellular Staining Perm Wash Buffer (BioLegend). Cells were then incubated with 1μg/ml CR3009 SARS-CoV-2 cross-reactive antibody (a kind gift from Dr. Laura McCoy) in permeabilization buffer for 30 min at room temperature, washed once and incubated with secondary Alexa Fluor 488-Donkey-anti-Human IgG (Jackson Labs). All samples were acquired on a BD Fortessa X20 using BD FACSDiva software. Data was analyzed using FlowJo v10 (Tree Star).

